# Coalescent inferences in conservation genetics: should the exception become the rule?

**DOI:** 10.1101/054528

**Authors:** Valeria Montano

## Abstract

Genetic estimates of effective population size (*N_e_*) are an established means to develop informed conservation policies. Another key goal to pursue the conservation of endangered species is keeping the connectivity across fragmented environments, to which genetic inferences of gene flow and dispersal greatly contribute. Most current statistical tools for estimating such population demographic parameters are based on Kingman's coalescent (KC). However, KC is inappropriate for taxa displaying skewed reproductive variance, a property widely observed in natural species. Coalescent models that consider skewed reproductive success-called multiple merger coalescent (MMCs)-have been shown to substantially improve estimates of Ne when the distribution of offspring per capita is highly skewed. MMCs predictions of standard population genetic parameters, including the rate of loss of genetic variation and the fixation probability of strongly selected alleles, substantially depart from KC predictions. These extended models also allow studying gene genealogies in a spatial continuum, providing a novel theoretical framework to investigate spatial connectivity. Therefore, development of statistical tools based on MMC's should substantially improve estimates of population demographic parameters with major conservation implications.

## Recent developments in coalescent theory

Estimates of effective population size (*N_e_*),defined by Wright as the number of reproducing lineages in an idealized population [1], are among the parameters used by the International Union for the Conservation of Nature (IUCN) to classify endangered species and to identify the minimum viable population size preventing extinction [2–3]. It has been suggested that IUCN thresholds of Ne recommended to avoid inbreeding depression and maintain evolutionary potential should be revised, as theoretical predictions often fail to match empirical observations [3]. However, a theoretical revision of *N_e_* thresholds will be ineffective to improve conservation recommendations if it is based on inappropriate evolutionary models.

Most methods applied in molecular ecology to infer demographic parameters from genetic data (e.g., Beast, Splatche, Ima, δaδi, FastSimcoal2, [4–8]) rely on Kingman's coalescent (KC; [9]) or its forward dual, the Wright-Fisher model (WF; [10]). Although KC has proven robust to violations of most of its assumptions, it drastically fails to approximate the genealogies of species with high reproductive skew [11], whereby few individuals contribute most of the offspring to the next generation (Sweepstakes Reproductive Success, or SRS [12]). Skewed disribution of per-capita reproductive success is widely observed among both marine and terrestrial species, from plants to parasites, but also among social birds and mammals [13]. SRS generally characterizes clonally reproducing organisms as much as species with high fecundity and low investment in parental care and thus applies to many endangered species, for instance amphibians and commercial fish.

Moreover, skewed individual reproductive success is not only due to intrinsic reproductive properties of a species, but can happen during strong population bottlenecks where only few individuals survive (e.g., a virus infecting a new host), during rapid population expansions [14], and during non-neutral processes, such as the appearance of a strongly beneficial allele which can drag a genome to replace an important fraction of the population within a few generations [15] (Figure 1).

**Figure 1.**
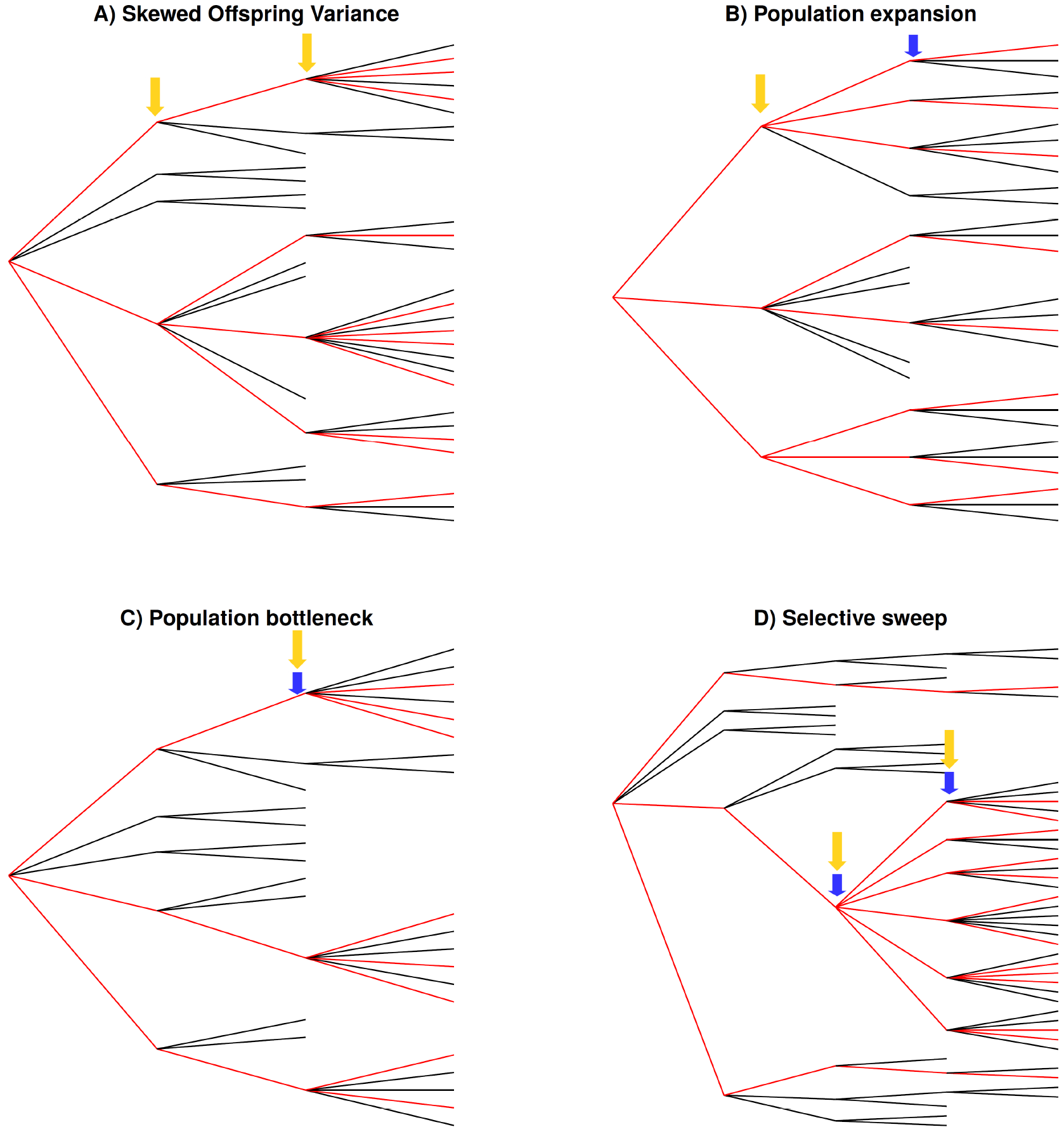
Examples of haploid genealogies presenting skewed reproductive success in forward and thus multiple merging in backward. Red edges indicate the sampled lineages. The yellow arrows represent the generation at which multiple merges occur and the blue arrows represent the generation at which the demographic event occurs. In A) SRS always leads to skewed offspring variance and thus multiple mergers can be observed at each generation, even when population size remains constant. In B) population expansion happens at the last generation with low reproductive variance and number of pre-capita offspring, hence the multiple mergers take place at the previous generation; in C) the population bottleneck and the multiple merging events occur at the same generation. In D) a selective sweep drags one genome to replace part of the population, thus the demographic event and the multiple merges co-occur.

KC model neglects the probability of more than two lineages to merge at each coalescent event, but when the offspring of a few individuals replaces a large fraction of the population at each reproductive event, the probability of multiple lineages merging in backward time becomes high. Hence, under skewed reproductive success, KC forces lineages involved in multiple and/or simultaneous merges to coalesce pairwise producing genealogical trees with misleading branch lengths and shape [11,14,15]. KC is a limit case of more complex coalescent processes, called multiple merger coalescences (MMCs), addressed in several recent studies, e.g., [11,12,14–18] and excellently reviewed in [18]. MMCs cover comprehensive scenarios, spanning from multiple lineages merging into one at each coalescent event (Λ-coalescent and its limit cases-β coalescent and Bolthausen-Sznitman coalescent [18]) to simultaneous multiple merging of multiple lineages at each coalescent event (Ξ-coalescent [18]). In MMC models, time-dependent changes in allele frequencies depart from KC predictions, consequently, probability of and time to fixation of both neutral and beneficial alleles, and, thus, the expected number of segregating sites dramatically change [19,20]. All of these measures are important to evaluate the health status of endangered species and their potential for adaptation to challenging environments [3].

When reproduction is highly skewed, few lineages substantially contribute to the next generation which means that the value of Ne, expressed by the parameter θ (2Neμ), is expected to be very low. However, under MMCs, alleles can persist at the same frequency for longer time than under KC before changing state, implying a reduced probability of loss or fixation for very low or high frequency alleles, respectively [19,20]. In contrast, when offspring variance and Ne are small, alleles at low frequencies are more likely to be lost by drift. Hence, under MMCs, the number of segregating sites and the number of singletons are predicted and empirically observed to assume close values, while under KC predicted number of singletons is usually much lower than number of segregating sites [11,16–18,21]. As a consequence, new beneficial mutations also show a higher chance to get lost under KC than under MMCs [19,20]. When few individuals contribute most of the offspring to the next generation, the frequency of few genotypes can increase substantially more than predicted by neutral KC. We can think of this scenario in terms of single lineages' rapid expansion, from which it follows that a high number of singletons can appear as the local genealogies become star-like. However, this scenario does not imply an expansion of the population size which can remain constant.

These differences between the KC and MMCs predictions explain two important results. First, MMCs estimates of Ne in marine species point to much lower values than KC estimates. In [11], the value of θ calculated for a population of oysters is 50 under KC and 0.031 under MMCs. From a conservation perspective, this result implies that high genetic variability can be generated by a very low number of lineages and thus an actual population might decline substantially without evident loss of genetic variation. At the same time, the ability of few individuals to quickly regenerate considerable genetic variation and the chance of new beneficial mutations to persist might result in high potential for rapid adaptation. Second, under MMCs and constant population size, a low θ value can recover both the observed number of segregating sites and singletons, while KC estimates fail to do so [11,21]. Therefore, conclusions pointing to population expansion based on excess of singletons-negative values of Tajima's D-should be carefully evaluated in molecular ecology studies.

## Spatial connectivity and continuous space evolution

Another theoretical advance of MMCs is the possibility to model continuous space evolution overcoming historical limitations. Indeed, models based on KC fail to control local population growth in continuous space, with the consequence that parts of the space grow unlimitedly and others become completely empty (a dynamic known as pain in the torus; [22,23]). As maintaining connectivity across habitats is indicated as a conservation priority [24], approaches to estimate connectivity in continuous landscapes based on circuit theory were developed as alternative to coalescent-based models [24,25]. Explicit spatial coalescent simulators based on KC (e.g,[5]) are still hampered by the use of discrete units which force coalescent events in non-contiguous populations [25] thus limiting their usefulness compared to alternative approaches [24,25]. In species with long distance dispersal ability and skewed reproductive success, local populations show low values of Ne associated to higher pairwise FST between closer than more distant populations [26]. This pattern can be explained by local bottlenecks due to few individuals reproducing and long distance dispersal events.

A forward model based on extinction-recolonization events (Λ-Fleming-Viot) allows to model evolution in spatial continuum using stochastic regulation of local size by randomly drawing the number of individuals destined for extinction (extinction event) and the number that will repopulate the same area from local or external parental lineages (recolonization event) [27,28]. The multiple merging spatial-Λ-coalescent is the backward dual of the forward Λ-Fleming-Viot processes [27,28]. Indeed, when lineages disappear backwardly during a recolonization event, multiple lineages will merge into the same or more parental individuals depending on how many parental lineages are responsible for the recolonization. When a parental lineage immigrates into a new area, the position of the descendant coalescing lineage will be spatially tracked back to a different part of the lattice corresponding to the origin of the parental lineage, such that the coalescing lineage is said to “jump” [27]. Allowing for local bottlenecks and long distance jumps, the spatial-Λ-coalescent can recover both small local *Ne* and long-distance correlated genealogies deriving from long distance dispersal events [27,28]. Without needing to assume discrete demes or homogeneous population distribution, this new framework has been shown to predict very well local and global N_e_ values when classic F_ST_ measures otherwise largely uncorrelate to observed values [26–29].

## Available statistical tools based on MMCs

Given the wide relevance of MMCs models to describe the demographic histories of natural populations (e.g., SRS, bottlenecks, expansions, positive selection), it is important to compare the fit of KC versus MMCs to describe a population demographic history, before a parameter of interest is estimated from empirical genetic data. While in species with highly skewed reproductive success MMCs can be assumed to outperform KC, in less trivial cases, e.g., human rapid population expansion [14], a model comparison is needed to accept or reject KC.

At the state of the art, some MMCs maximum likelihood estimators have been developed and are available to infer the effective population size and skewness of the offspring distribution of marine species [11,25,30], such as Metagenetree [17] (Table 1). A recent software based on spatial-Λ- coalescent (*phyrex*) by [29] estimates global *Ne* values in continuous space as an alternative to classic *F_ST_* estimates. Moreover, two MMCs simulators are currently available: algorithms by Kelleher et al for continuous space evolution [29] and Hybrid-Lambda for species evolution [31], which could be used to fit evolutionary hypotheses to observations using simulation approaches (Table 1). Indeed, Joseph et al 2016 [32] developed an ABC pipeline based on the simulator presented in [29] (Table 1). At the same time, empirical conservation biologists will benefit from being aware of the biological relevance of MMCs and when and why they should be applied.

**Table 1.** Available statistical tools based on MMC models.

| MMC tools |  |  |  |  |  |
| --- | --- | --- | --- | --- | --- |
| *Name* | *Type* | *Model* | *Spatially explicit* | *Reference* | *Source* |
| Eldon & Wakeley | Estimator | Λ-coalescent | No | Eldon and Wakeley 2006 | Available under request to authors |
| Metagenetree | Estimator | Λ-coalescent | No | Birkner et al 2011 | http://metagenetree.sourceforge.net/ |
| Phyrex | Estimator | Spatial-Λ-coalescent | Yes | Guindon et al 2016 | https://github.com/stephaneguindon/phyml |
| Hybrid-Lambda | Simulator | B and Λ-coalescent | No | Zhu et al 2015 | https://github.com/hybridLambda/hybrid-Lambda |
| ABC-Discsim | Simulator and Estimator | Spatial-Λ-coalescent | Yes | Kelleher et al 2014; Joseph et al 2016 | https://github.com/tyjo/ABC-Discsim |

## Acknowledgements

I am grateful to Mauricio Gonzalez-Forero, Jeffrey Jensen, Sebastian Matuszewski, Stefan Laurent, Oscar Gaggiotti, Chiara Batini and two anonymous reviewers for helpful comments.

